# Transcriptome-wide Analyses of Adipose Tissue in Outbred Rats Reveal Genetic Regulatory Mechanisms Relevant for Human Obesity

**DOI:** 10.1101/2022.03.24.485632

**Authors:** Wesley L. Crouse, Swapan K Das, Thu Le, Greg Keele, Katie Holl, Osborne Seshie, Ann L Craddock, Neeraj K. Sharma, Mary Comeau, Carl D Langefeld, Greg Hawkins, Richard Mott, William Valdar, Leah C Solberg Woods

**Affiliations:** University of North Carolina at Chapel Hill, Department of Genetics, Chapel Hill, NC, USA; Wake Forest University School of Medicine, Department of Internal Medicine, Winston Salem, NC, USA; University College London, Department of Genetics, Evolution and Environment, Division of Biosciences, London, UK; Jackson Laboratories, Roux Center for Genomics and Computational Biology, Bar Harbor, ME, USA; Medical College of Wisconsin, Department of Pediatrics, Milwaukee, WI, USA; Wake Forest University School of Medicine, Department of Biochemistry, Winston Salem, NC, USA; Wake Forest University School of Medicine, Department of Biostatistics and Data Sciences, Winston Salem, NC, USA

**Keywords:** RNA-seq, WGCNA, mediation analysis, network analysis, adipose tissue Short running title: Adipose transcriptome networks in rat and human

## Abstract

Transcriptomic analysis in metabolically active tissues allows a systems genetics approach to identify causal genes and networks involved in metabolic disease. Outbred heterogeneous stock (HS) rats are used for genetic mapping of complex traits, but to-date, a systems genetics analysis of metabolic tissues has not been done. We investigated whether adiposity-associated genes and gene co-expression networks in outbred heterogeneous stock (HS) rats overlap those found in humans. We analyzed RNAseq data from adipose tissue of 415 male HS rats, correlated these transcripts with body weight (BW) and compared transcriptome signatures to two human cohorts: the African American Genetics of Metabolism and Expression and Metabolic Syndrome in Men. We used weighted gene co-expression network analysis to identify adiposity-associated gene networks and mediation analysis to identify genes under genetic control whose expression drives adiposity. We identified 554 orthologous “consensus genes” whose expression correlates with BW in the rat and with body mass index (BMI) in both human cohorts. Consensus genes fell within eight co-expressed networks and were enriched for genes involved in immune system function, cell growth, extracellular matrix organization and lipid metabolic processes. We identified 19 consensus genes for which genetic variation may influence BW via their expression, including those involved in lipolysis (e.g., *Hcar1)*, inflammation (e.g., *Rgs1*), adipogenesis (e.g., *Tmem120b*) or no previously known role in obesity (e.g., *St14, Msa4a6*). Strong concordance between HS rat and human BW/BMI associated transcripts demonstrates translational utility of the rat model, while identification of novel genes expands our knowledge of the genetics underlying obesity.

## INTRODUCTION

Obesity and overweight are major risk factors for multiple diseases including cardiovascular disease, type 2 diabetes and cancer (1). There has been a steady increase in prevalence of overweight and obesity since the 1970’s (2), and by 2030, approximately half of the adults in the U.S. are expected to be obese (3). Obesity is caused by genetic and environmental factors with genetic factors accounting for up to 70% of the population variance (4). To date, human genome wide association studies (GWAS) have identified several hundred genomic loci for body mass index (BMI) and wait-hip-ratio (WHR), but these loci explain only 6% of the heritable variation (5, 6), indicating much is left to be found.

Animal models are routinely used for mechanistic understanding of metabolic disease including obesity. We have previously used association analysis in outbred heterogeneous stock (HS) rats to identify both novel and known genes involved in adiposity (7, 8) and other metabolic traits (9–11). HS rats were created by combining eight inbred founder strains and maintaining the colony in a way that minimizes inbreeding (12), thereby mimicking the genetic diversity seen in humans. The chromosomes of each HS animal are fine-grained mosaics of the founder haplotypes, allowing genetic fine-mapping to Mb-sized intervals (13). However, evidence that HS rats recapitulate human obesity at a molecular level and on a genome-wide scale is lacking.

Transcriptomic analysis in metabolically active tissues can identify both causal and reactive genes and networks involved in metabolic disease. Adipose tissue plays a central role in metabolic health (14). Although human BMI GWAS genes show strong enrichment in the central nervous system (15), adipose tissue-expressed genes are also enriched in human GWAS for BMI (16) and to a greater extent for WHR (17, 18). In addition, adipose tissue function is disrupted under high fat diet (HFD) and/or obese conditions (19), making it a highly relevant tissue for the study of obesity. Although transcriptome analysis has been conducted in adipose tissue of recombinant inbred mouse lines (20), this work has not previously been conducted for HS rats. Thus, we investigated whether adiposity-associated genes and gene co-expression networks in outbred heterogeneous stock (HS) rats overlap with those found in humans. We determined the concordance of body weight (BW)/BMI-associated adipose tissue transcripts between HS rats and two human cohorts, namely the extensively gluco-metabolically phenotyped human participants in the African American Genetics of Metabolism and Expression (AAGMEx) (21) and Metabolic Syndrome in Men (METSIM) (22).

Importantly, networks can include both genetically driven causal genes and those that are reactive to the phenotype. By leveraging genetic information and performing mediation analysis in HS rats, we further identified a subset of rat/human consensus genes which may be causal for obesity. Thus, this work not only sheds light on cross-species gene networks involved in BW/BMI, but also employs a complementary approach to human GWAS for identifying novel gene regulators of adiposity.

## RESEARCH DESIGN AND METHODS

### Animals

The HS rat colony was initiated by the NIH in 1984 by breeding together the following eight inbred founder strains: ACI/N, BN/SsN, BUF/N, F344/N, M520/N, MR/N, WKY/N and WN/N, and maintaining the colony in a way that minimizes inbreeding (12). The HS colony had been maintained at the Medical College of Wisconsin (MCW; NMcwi:HS; RGD_2314009) from 2006 – 2017, after which time a colony was set up at Wake Forest School of Medicine (WFSM; NMcwiWFsm:HS; RGD_13673907) (13). Animals for the current study came from HS rats maintained at the MCW colony and were collected from 2006 to 2011. Animals were housed 2 per cage in micro-isolation cages in a conventional facility using autoclaved bedding (sani-chips from PJ Murphy). They were given ad libitum access to autoclaved Teklad 5010 diet (Harlan Laboratories) and were provided reverse osmosis water chlorinated to 2-3 ppm. 1144 male HS rats went through the phenotyping protocol below, running approximately 12 animals/batch with a new batch run each week. Retroperitoneal adipose tissue (RetroFat) from 415 of these rats was used for transcriptomic analysis, as described below.

### HS rat phenotyping protocol

As previously described (10), at 16 weeks of age, body weight (BW) was measured after an overnight fast (∼16 hours), after which time rats underwent an intra-peritoneal glucose tolerance test (IPGTT). The IPGTT was conducted under the hood in the animal room and was conducted in the morning from ∼9am – 12pm. Blood and plasma were collected at 0, 15, 30, 60 and 90 minutes after a 1g/kg BW glucose injection. Blood glucose was measured using the Ascensia Elite system (Bayer, Elkhart, IN) and plasma insulin levels were determined using an ultrasensitive ELISA kit (Alpco Diagnostics, Salem, NH). Using blood glucose and plasma insulin levels, we calculated area under the curve for glucose (glucose_AUC) and insulin (insulin_AUC) during the IPGTT as previously described (10). At 17 weeks of age rats were euthanized after an overnight fast. At the time of euthanasia, BW and body length (BL) were measured. Rats were then euthanized by decapitation and trunk blood was collected. Fasting cholesterol and triglycerides were determined from fasting serum on an ACE Alera autoanalyzer using an enzymatic method for detection. We collected and snap froze several tissues including RetroFat and epididymal fat (EpiFat) pads for subsequent expression analysis. We also stored whole pancreas in acid ethanol for subsequent determination of whole pancreas insulin content. All protocols were approved by the IACUC committee at MCW. Phenotyping data has been deposited in RGD (www.rgd.mcw.edu; RGD: 151665312).

### HS rat Genotyping and Imputation

We extracted DNA from tail tissue of all 1144 samples using a phenol-chloroform extraction. All samples were genotyped by obtaining low coverage whole genome sequence (0.24x, performed by Beijing Genomics Institute (BGI)) followed by imputation by STITCH, as previously described (23). Eight founders with high coverage sequence (24-28X) (24) were used as the haplotype reference panel in STITCH, yielding imputed single nucleotide polymorphism (SNP) calls at approximately 4.7M sites in the HS rats. Quality control for SNP calls included retaining SNP calls with imputation info score > 0.4 and Hardy-Weinberg Equilibrium p-value > 10^−6^, and removing SNPs with high linkage disequilibrium R^2^ > 0.95. After quality control, a tagging set of more than 122,000 SNPs were used for further analyses.

### HS rat RNAseq analysis

We used Trizol to extract RNA from RetroFat of 415 HS rats. RNA quality was confirmed using a Bioanalyzer. Illumina kits were used to create library preps and RNA-seq was run on the Illumina HiSeq 2500 by the Wake Forest University School of Medicine Genomics Core, obtaining 75-bp single end reads. STAR (25) was used to align reads to the rat genome reference 6.1 and DESeq2 (26) was used to compute gene level expression abundance. We excluded very low expressed genes, where the average number of reads per sample < 1. Read coverage for each remaining gene was then normalized to account for gene length. A total of 18,357 genes from adipose tissue were used for further analyses. RNAseq data has been submitted to Gene Expression Omnibus (GEO); id #GSE196572.

### Quality Control and Pre-processing of the HS rat RNAseq data

We first performed exploratory principal components analysis and determined the mean-variance relationship between all genes in the dataset. Based on this analysis, we visually identified 28 genes that were considerably more variable than other genes with similar levels of expression (mean expression < 100 and variance > 700; see **Fig. S1**). These 28 genes were significantly enriched for Gene Ontology (GO) terms related to muscle cells (see **Table S1** and ‘Enrichment analysis’ section below) and also included the gene MPZ, which is a marker for nerve cells (27). The high variance of these genes suggest tissue heterogeneity across samples, with varying proportions of muscle and nerve cells mixed in with adipose tissue. We performed additional pre-processing steps on the phenotypes and gene expression data prior to further analysis, as described in detail below.

**Table 1.** Adipose tissue WGCNA modules associated with body weight and other metabolic traits

| Module Color<br>[Total gene #<br>(# of genes most<br>positively/inversely<br>correlated with<br>BWg at \|r\|≥0.2)] | Biological pathway<br>Top GO.BP*<br>(FDR-P) | Body Weight<br>(p-value) † | RetroFat<br>(p-value) † | EpiFat<br>(p-value) † | Fasting<br>Insulin<br>(p-value) † | Fasting<br>Cholesterol<br>(p-value) † | Fasting<br>Triglycerides<br>(p-value) † | # of consensus gene<br>enriched in module<br>(percent; padj for<br>enrichment) |
| --- | --- | --- | --- | --- | --- | --- | --- | --- |
| **Positive Associations** |  |  |  |  |  |  |  |  |
| Light Green<br>[141 (60/10)] | Regulation of pathway-restricted<br>SMAD protein phosphorylation<br>(0.008) | **0.35**<br>(1.6x10<sup>-13</sup>) | **0.54**<br>(4.8x10<sup>-33</sup>) | **0.46**<br>(9.5x10<sup>-23</sup>) | **0.34**<br>(1.7x10<sup>-12</sup>) | **0.31**<br>(6.4x10<sup>-11</sup>) | **0.35**<br>(1.3x10<sup>-13</sup>) | 22<br>(15.6%; 1.9x10<sup>-09</sup>) |
| LightYellow<br>[139 (62/7)] | Extracellular matrix organization<br>(4.2x10<sup>-4</sup>) | **0.33**<br>(6.9x10<sup>-12</sup>) | **0.15**<br>(0.002) | **0.19**<br>(1.2x10<sup>-04</sup>) | -0.03<br>(NS) | 0.01<br>(NS) | -0.03<br>(NS) | 11<br>(13.7%; 2.4x10<sup>-07</sup>) |
| LighyCyan<br>[148 (27/0)] | Cell Cycle Process<br>(2.8x10<sup>-48</sup>) | **0.23**<br>(1.5x10<sup>-06</sup>) | **0.14**<br>(0.005) | **0.17**<br>(7.3x10<sup>-04</sup>) | 0.05<br>(NS) | 0.02<br>(NS) | -0.02<br>(NS) | 8<br>(5.4%; 0.266) |
| Dark Red<br>[107 (23/1)] | Complement Activation<br>(2.7x10<sup>-08</sup>) | **0.22**<br>(3.8x10<sup>-06</sup>) | **0.11**<br>(0.028) | **0.11**<br>(0.029) | **0.10**<br>(0.053) | 0.07<br>(NS) | -0.05<br>(NS) | 21<br>(19.6%; 7.2x10<sup>-11</sup>) |
| Brown<br>[1138 (109/49)] | Immune system process (2.8x10<sup>-48</sup>) | **0.20**<br>(3.2x10<sup>-05</sup>) | **0.14**<br>(0.005) | **0.18**<br>(2.6x10<sup>-04</sup>) | 0.036<br>(NS) | **0.12**<br>(0.011) | 0.04<br>(NS) | 113<br>(9.9%; 7.9x10<sup>-29</sup>) |
| Magenta<br>[472 (26/19)] | Golgi vesicle transport<br>(0.046) | **0.18**<br>(1.9x10<sup>-04</sup>) | **0.26**<br>(7.5x10<sup>-09</sup>) | **0.17**<br>(3.5x10<sup>-04</sup>) | **0.22**<br>(6.4x10<sup>-06</sup>) | **0.12**<br>(0.017) | **0.11**<br>(0.024) | 16<br>(3.4%; 0.962) |
| DarkGreen<br>[98 (6/0)] | Response to Virus<br>(5.3x10<sup>-35</sup>) | **0.16**<br>(0.001) | -0.04<br>(NS) | -0.02<br>(NS) | -0.06<br>(NS) | -0.00<br>(NS) | **-0.13**<br>(0.008) | 4<br>(4.0%; 0.962) |
| Purple<br>[445 (20/4)] | Extracellular matrix organization<br>(1.9x10<sup>-12</sup>) | **0.13**<br>(0.007) | 0.02<br>(NS) | 0.02<br>(NS) | -0.08<br>(NS) | 0.03<br>(NS) | -0.07<br>(NS) | 10<br>(2.2%; 1) |
| Grey60<br>[142 (7/1)] | tRNA aminoacylation for protein<br>translation<br>(0.007) | **0.11**<br>(0.026) | **0.33**<br>(9.8x10<sup>-12</sup>) | **0.20**<br>(2.7x10<sup>-05</sup>) | **0.27**<br>(2.00x10<sup>-08</sup>) | **0.10**<br>(0.038) | **.20**<br>(3.5x10<sup>-05</sup>) | 4<br>(2.8%; 1) |
| **Negative Associations** |  |  |  |  |  |  |  |  |
| Green<br>[668 (71/111)] | Cytokine Production<br>(4.7x10<sup>-05</sup>) | **-0.31**<br>(2.2x10<sup>-10</sup>) | **-0.23**<br>(1.4x10<sup>-06</sup>) | **-0.23**<br>(3.3x10<sup>-06</sup>) | **-0.13**<br>(0.008) | **-0.16**<br>(0.001) | -0.03<br>(NS) | 78<br>(11.6%; 4.6x10<sup>-24</sup>) |
| DarkTurquoise<br>[90 (3/1)] | Circadian Rhythm<br>(0.007) | **-0.21**<br>(1.3x10<sup>-05</sup>) | -0.06<br>(NS) | -0.04<br>(NS) | **0.15**<br>(0.002) | 0.04<br>(NS) | 0.09<br>(NS) | 9<br>(10%; 0.006) |
| Blue<br>[1247 (23/71)] | Mitochondrion organization<br>(3.4x10<sup>-19</sup>) | **-0.15**<br>(0.002) | -0.04<br>(NS) | **-0.13**<br>(0.010) | 0.09<br>(NS) | -0.07<br>(NS) | -0.02<br>(NS) | 66<br>(5.2%; 2.8x10<sup>-05</sup>) |
| Cyan<br>[203 (1/20)] | Regulation of Neurotransmitter<br>Levels<br>(0.004),<br>Lipid Metabolic Process<br>(0.008) | **-0.12**<br>(0.014) | **-0.29**<br>(2.4x10<sup>-09</sup>) | **-0.30**<br>(6.3x10<sup>-10</sup>) | **-0.23**<br>(1.3x10<sup>-06</sup>) | **-0.29**<br>(3.2x10<sup>-09</sup>) | **-0.29**<br>(1.2x10<sup>-09</sup>) | 13<br>(6.4%; 0.033) |
| Yellow<br>[750 (11/18)] | Nucleic Acid Metabolic Process<br>(4.7x10<sup>-10</sup>) | **-0.10**<br>(0.046) | **-0.17**<br>(7.0x10<sup>-04</sup>) | **-0.14**<br>(0.004) | **-0.14**<br>(0.003) | -0.01<br>(NS) | **-0.11**<br>(0.023) | 20<br>(2.7%; 1) |

\*Top most enriched gene ontology terms among gene members of each module are shown; †correlation and p-value of module Eigengene with each trait are shown. Those that reach statistical significance are in bold. Modules enriched for consensus genes are highlighted in grey.

#### Phenotypes

Each of the phenotypes were pre-processed by removing covariates and transforming the residuals to follow an approximately normal distribution. The traits EpiFat and fasting cholesterol and triglyceride were log transformed based on a Box-cox procedure, while all other phenotypes were transformed using a rank inverse normal (RINT) transformation. We then adjusted the transformed phenotypes for phenotype-specific experimental covariates using linear regression with random covariate effects (see **Table S2**). This is given by

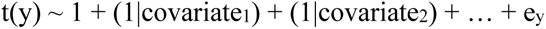

where t(y) is a transformed phenotype, 1 denotes the intercept, (1|covariatei) denotes random effects for the i-th covariate, and ey is the residual. We used the residuals of this regression, denoted y’=ey, in further analyses.

#### Adipose Gene Expression

We transformed adipose gene expression using RINT. In order to adjust the adipose expression for tissue heterogeneity, we created proxy variables for nerve and muscle content. The proxy variable for nerve content was the transformed expression of the MPZ gene. The proxy variable for muscle content was the first principal component of the transformed expression of the other 27 genes identified during exploratory analysis. This first principal component explained 57.5% of the variation in expression for this subset of muscle-related genes. These proxy variables were negatively correlated with several weight-related phenotypes, and most strongly with Retrofat, indicating a higher amount of muscle contamination in rats with smaller fat pads, likely due to dissection technique (see **Fig. S2**).

We then adjusted the transformed gene expression for tissue composition and experimental batch using linear regression with fixed tissue-proxy effects and random batch effects. This is given by

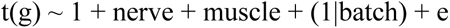

where t(g) is transformed adipose gene expression, ‘nerve’ and ‘muscle’ denote fixed effects for the nerve and muscle proxy variables, (1|batch) denotes random effects for RNAseq batches (e.g., those samples that were sequenced together), and e is the normally-distributed residual error. We used the residuals of this regression, denoted g’=e, in further analyses.

### Associations with phenotypes

After quality control, and for each transformed phenotype, we used linear regression to identify genes with significantly associated expression in adipose tissue; significance was defined as having a false discovery rate (FDR) of less than 1%, with FDR calculated using the Benjamini and Hochberg procedure (B-H) (28).

### Weighted Gene Co-expression Network Analysis (WGCNA)

We identified modules of co-expressed genes that correlated with the metabolic phenotypes (29). This was done using the WGCNA R package (30) using the recommended approach for automatic network construction and module detection in the WGCNA tutorial. We assumed signed correlation networks and computed bi-weight mid-correlations between adipose tissue genes. A threshold of 0.9 was used for the scale free topology index to set the soft thresholding power, which was set to 5 (see scale free topology and mean connectivity in **Fig. S3**). Recommendations from the tutorial were used for settings related to dendrogram cutting and module merging. Module eigengenes (first principal component of genes included in the module) were then correlated with each phenotype using Spearman correlation. The function of each module was characterized using gene ontology (GO) enrichment. Module membership and intermodular connectivity (IMConn) was computed for all genes using the adjacency matrix from the WGCNA analysis in order to identify highly-connected hub genes within each module. We defined hub genes within each module using a threshold of IMConn > mean + 2 standard deviations. Gene networks were visualized for selected modules using Cytoscape. For visualization in Cytoscape, we applied a minimum adjacency threshold of 0.03 for including edges between genes, and we removed genes that were not connected to the main module network.

### Gene Ontology (GO) Enrichment and Pathway analysis

GO enrichment analysis was performed on genes significantly associated with each phenotype and modules of co-expressed genes. We used the ‘anRichment’ R package (https://horvath.genetics.ucla.edu/html/CoexpressionNetwork/GeneAnnotation/). This performs a Fisher’s exact test for enrichment, and we assumed that the background for this test was the intersection of all genes analyzed and all genes with GO annotations. FDR was controlled at 5% using B-H. For tests of associated genes, FDR was controlled within phenotype, and for tests of modules, FDR was controlled across modules. We also evaluated enrichment of differentially expressed genes for KEGG pathways (using the Database for Annotation, Visualization and Integrated Discovery/DAVID; https://david.ncifcrf.gov/) and for canonical pathways in Ingenuity Pathway Knowledgebase (using Ingenuity Pathway Analysis, IPA).

### AAGMEx and METSIM human cohorts

In order to identify consensus genes associated with adiposity in both rats and humans, we compared the adipose gene expression data in HS rats with adipose expression from two separate human cohorts: AAGMEx and METSIM. The AAGMEx cohort consisted of 256 healthy, self-reported African American men and women residing in North Carolina, aged 18-60 years, with a body mass index (BMI) between 18 and 42 kg/m^2,^. Participants were unrelated and non-diabetic. Clinical, anthropometric, physiological characteristics, and detail of genomic data processing of the AAGMEx cohort have been described previously (21, 31). Subcutaneous adipose tissue biopsies were collected by Bergstrom needle from participants after an overnight fast. Genome-wide expression data of subcutaneous adipose tissue biopsies (submitted to GEO; id #GSE95674) in AAGMEx were generated using HumanHT-12 v4 Expression BeadChip (Illumina, San Diego, CA) whole-genome gene expression arrays, and Infinium HumanOmni5Exome-4 v1.1 DNA Analysis BeadChips (Illumina) were used for genotyping. Previously completed statistical analyses on BMI, adipose tissue gene expression, and genotype data were used for validation /replication of findings from rat in human. We also used previously published statistical analyses results on BMI and adipose tissue gene expression data from the European Ancestry individuals in the METSIM cohort which consisted of 770 male individuals from Finland, as previously described (METSIM, Finland; N=770) (32). In the METSIM cohort, subcutaneous adipose tissue expression data were generated by Affymetrix U219 arrays (GEO id # GSE70353).

### Identifying adipose tissue consensus genes associated with BW in rat and BMI in human

We filtered the 2,419 BW-associated genes from rat adipose tissue to include only genes with human orthologues (2,200 genes). For each gene, we checked if adipose tissue expression of its human orthologue was significantly associated (FDR = 1%) with BMI independently in both the AAGMEx and METSIM cohorts, with the same direction of effect. Specifically, to test for associations between BMI and expression level in the AAGMEx cohort, we computed a linear regression model using R(lm) software with the BMI (square root transformed) as the outcome and expression level (RMA normalized, batch corrected, and log2 transformed) as the predictor. Models included age, gender, and African ancestry proportion (admixture estimates were computed using the ADMIXTURE program (http://software.genetics.ucla.edu/admixture/index.html) as covariates. Similar regression analyses were conducted to evaluate the association between gene expression and cardio- metabolic-related traits, including BMI in up to 770 METSIM individuals. In METSIM, both BMI values and RMA-normalized (non-PEER-corrected) expression levels were adjusted for age before a rank inverse normal transformation. These transformed values were used for computing association between gene expression and BMI.

Upon identifying genes that were associated with BW in rat and BMI in human with the same direction of effect, we ran a chi-square test to determine if the number of consensus genes is higher than would be expected by chance. We then used a Fisher combined p-value with equal weights to rank these consensus genes by their significance in all three datasets. We also tested for enrichment of these genes in the WGCNA adipose modules using Fisher’s exact test in order to identify modules enriched for many consensus genes.

### Mediation analysis

Our consensus gene approach does not distinguish between genes that are either causal or reactive to BW. In order to assess evidence in favor of a causal relationship between consensus genes and BW, we leveraged SNP information in the rat dataset to identify cis-eQTL for consensus genes. We then tested if these cis-eQTL are associated with BW, as part of a formal mediation analysis based on (8, 33).

Typically, mediation analysis is performed by first finding a genetic association with a trait, and then testing potential mediators of this relationship. For example, a classic procedure for mediation analysis is given by Baron and Kenny (33), which in our context involves sequentially testing for the following relationships:

1. The genetic variant x is associated with the phenotype y’:

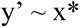
2. The genetic variant is associated with the mediator gene g’:

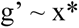
3. The mediator gene is associated with the phenotype in the presence of the genetic variant:

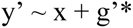
4. The genetic variant is not associated with the phenotype in the presence of the mediator gene:

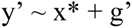

In these expressions, the (*) denotes which dependent variable is being tested for association. If the tests of association in Steps 1-3 are all significant, then there is evidence for partial mediation of the genetic effect through the gene. Additionally, if the test of association in Step 4 is not significant, then there is evidence for complete mediation.

Our application of mediation analysis is nonstandard, in that we already identified a set of consensus genes that are associated with BW/BMI, and we use mediation to establish a link between the genetics of these genes and our trait. This is different than a typical application of mediation analysis (e.g., (8, 34)), in which Step 1 is already satisfied, perhaps with genome-wide significance. Given that we already have a list of candidate genes to assess, and we are only interested in their local genetic variation, a genome-wide significance threshold for genetic association in either Step 1 or Step 2 is overly conservative. We also note that, prior to performing mediation analysis, we did not strictly test Step 3, but our approach for identifying consensus genes did test a similar hypothesis (y’ ∼ g’*). Given these considerations, we did not use the typical sequential approach as described by Baron and Kenny, and instead used the following procedure to assess evidence for mediation.

For each consensus gene, we first performed local cis-eQTL mapping of rat gene expression (Step 2) using all SNPs within a 1Mb window of the gene start or end. We selected the most significant SNP for each gene for further analyses. To assess eQTL significance, we applied a two-stage FDR approach to account for testing multiple SNPs per gene across multiple genes. First, we applied a B-H correction within gene across all SNPs. Then, we applied a second B-H correction to the adjusted p-value of the most significant SNP across all genes. SNPs were deemed significant eQTL if their two-stage adjusted p-values were significant (FDR = 5%). Then, for all consensus genes with significant eQTL, we tested Steps 1, 3, and 4 using nominal significance thresholds (i.e., no multiple testing correction). In the results, we report nominal p-values for Steps 1-4, for all genes satisfying Steps 1-3, which are considered partial mediators. We then used PhenomeXcan (http://apps.hakyimlab.org/phenomexcan/) to determine if mediation genes in rat exhibit causal gene-trait associations in human (35). Specifically, for each of the 19 genes, we identified gene-trait associations for fat mass related phenotypes (p < 0.01) for both subcutaneous and visceral (omental) adipose tissues.

## Data and Resource Availability

All supplemental figures and tables can be accessed using the following figshare link: https://figshare.com/s/9bdfdd07d80744382963. The datasets generated and analyzed during the current study are also available on figshare: https://figshare.com/articles/dataset/hs_rat_adipose_gene_expression_analysis_zip/16620583/2.

Phenotype data has been submitted to the Rat Genome Database (https://rgd.mcw.edu/; RGD: 151665312) and RNAseq data has been submitted to Gene Expression Omnibus GEO (id #GSE196572).

The HS rats used in the current study, now maintained at Wake Forest University School of Medicine (WFSM; NMcwiWFsm:HS) are available from the corresponding author upon reasonable request and on a cost-recovery basis.

## RESULTS

Analysis framework and key results, including the relationship between correlation analysis, WGCNA, enrichment in human cohorts and mediation analysis are shown in **Figure 1**.

**Figure 1.**
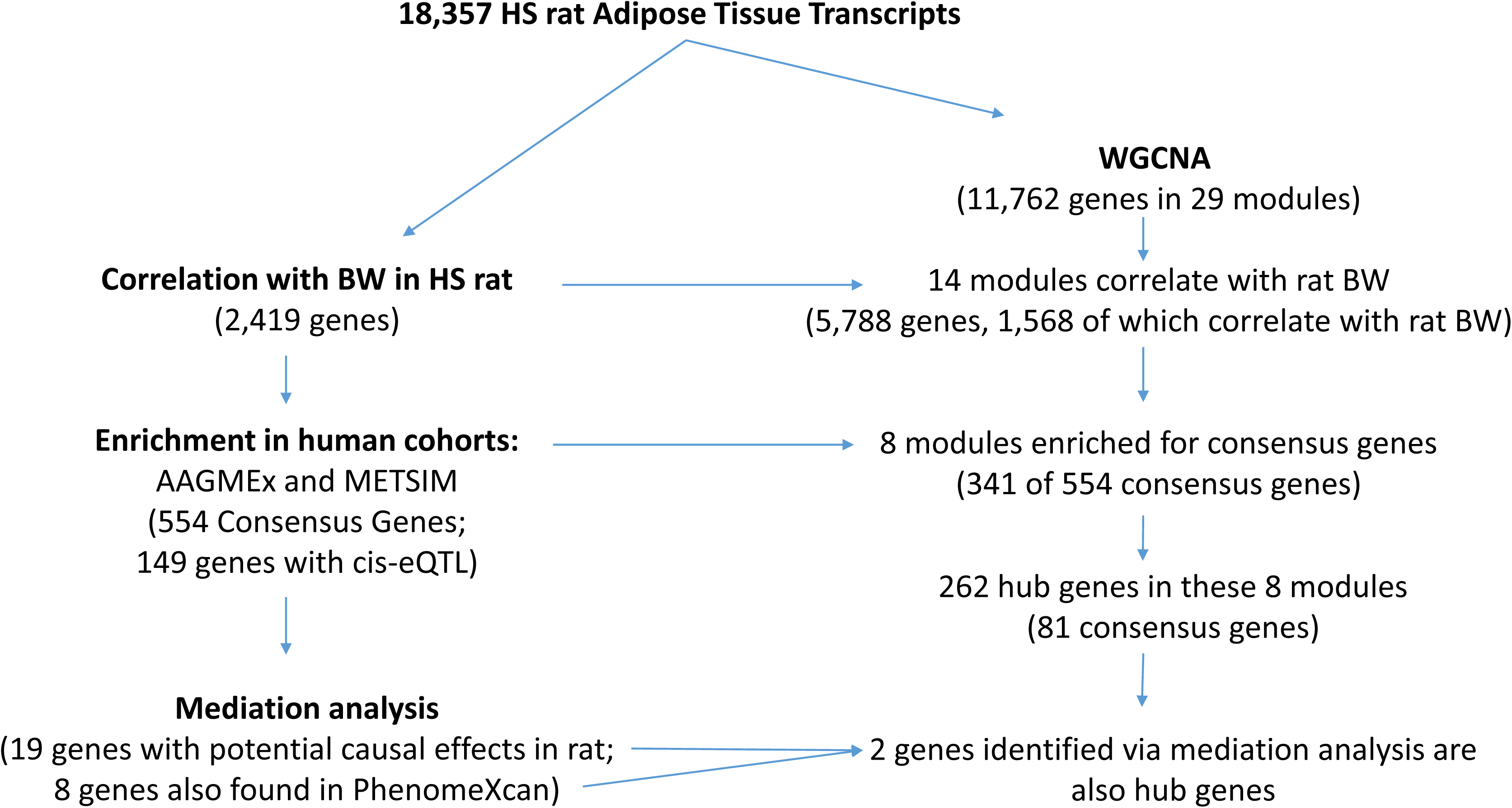
Analysis framework and key results. Figure depicts the connection between correlation analysis (adipose tissue transcript and HS rat body weight) and WGCNA networks. Correlation analysis was followed by identification of human/rat BMI/BW consensus genes and then mediation analysis. The number of genes or modules at each step are shown in parentheses. Horizontal arrows depict how and where correlation and consensus analysis feed into WGCNA modules.

### BW-associated transcripts in rat adipose tissue are enriched for relevant gene-ontology categories and biological pathways

We identified 2,419 genes with normalized expression levels significantly associated with rat BW after multiple testing correction (FDR = 1%), with 1,277 genes positively and 1,142 genes negatively associated. The most positively associated genes were *Cpa1* (beta = 0.499, p = 3.2 x 10-23)*, Bace2* (beta = 0.411, p = 1.4 x 10-14), and *Dclk1* (beta = 0.403, p = 5.1 x 10-14); the most negatively associated genes were *Fmo1* (beta = -0.411, p = 1.4 x 10-14)*, Acad8* (beta = -0.400, p = 7.5 x10-14) and *Cds1* (beta = -0.375, p = 5.8 x10-12) (**Table S3**). Of the associated genes, 1,665/2,419 (68.8%) were also associated with at least one other measured metabolic phenotype. This is not unexpected given that many of the metabolic phenotypes are correlated (**Fig. 2**). The 2,419 BW-associated transcripts were enriched for 106 GO terms at FDR of 5% (**Table S4**). Similar evaluation for the enrichment of KEGG pathway annotations identified 56 pathways, five of which survived multiple testing at FDR of 5%. These include: *receptor mediated phagocytosis, protein digestion and absorption, chemokine signaling pathway, protein processing in endoplasmic reticulum and aldosterone regulated sodium reabsorption* (**Table S5**) which are generally supported by canonical pathway enrichment analysis using IPA (**Table S6**).

**Figure 2.**
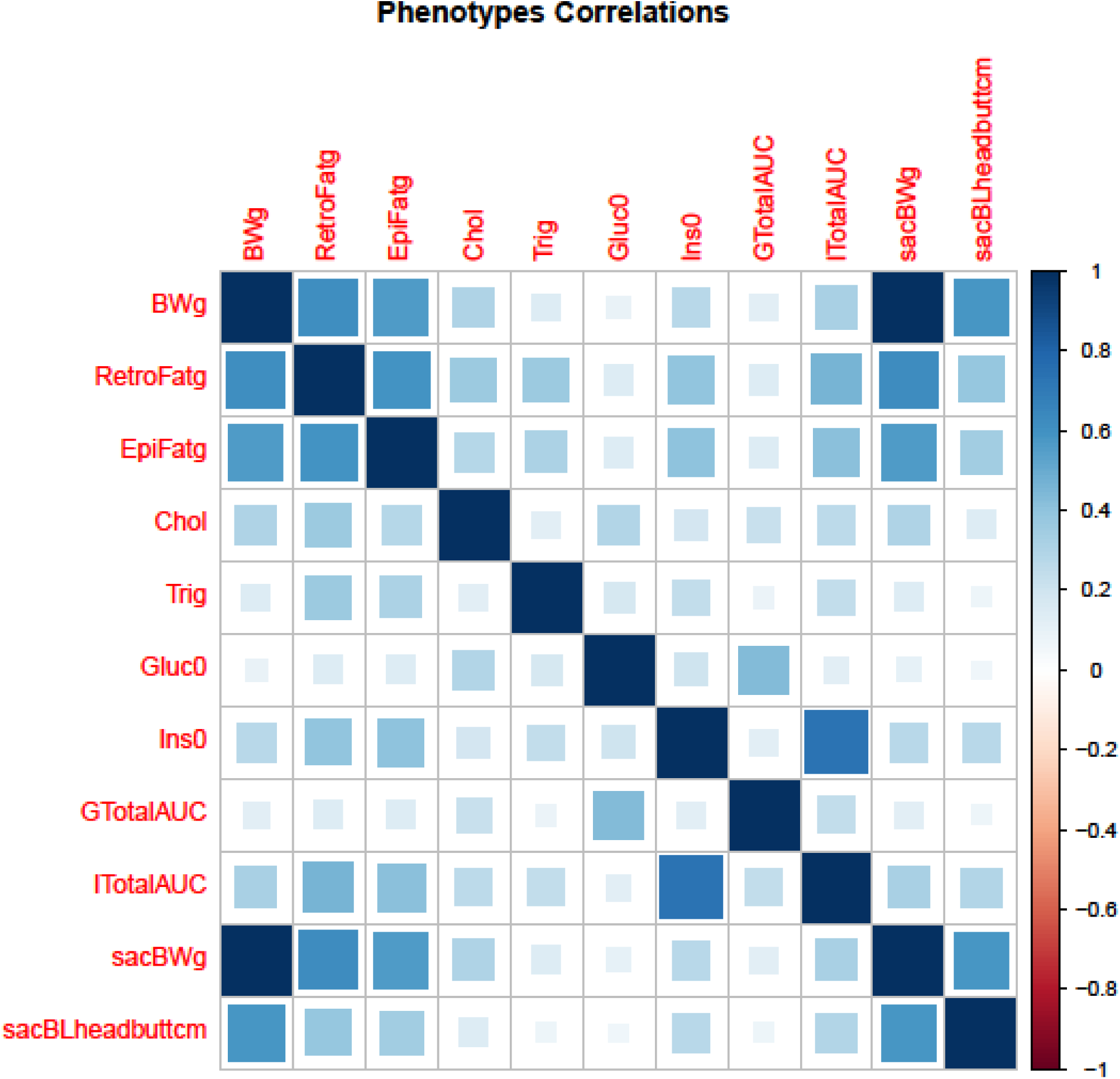
Correlations between metabolic phenotypes in HS rats. Pearson correlations between multiple metabolic phenotypes in 1144 HS male rats. Darker color indicate a higher correlation. Positive correlations are shown in blue. All metabolic phenotypes are positively correlated. Adiposity traits are highly correlated with each other and with insulin traits. BWg – bodyweight (g); BLcm – body length (cm); EpiFatg – epididymal fat pad weight (g); RetroFatg – retroperitoneal fat pad weight (g); Ins0 – fasting insulin (ng/ml); InsAUC – insulin area under the curve after glucose challenge; Gluc0 – fasting glucose (mg/dL); GlucAUC – glucose area under the curve after glucose challenge; Chol – fasting total cholesterol (mg/dL); Trig – fasting triglycerides (mg/DL).

### Gene co-expression analysis identifies biologically-relevant modules associated with BW in rat adipose tissue

Gene co-expression analysis using WGCNA assigned 11,762 of the 18,357 expressed transcripts into 29 modules or sub-networks (**Figs. 3 and 4**). Eigengenes of 14 of these modules were associated with BW (p=0.05) (**Table 1**). The modules most strongly positively associated with BW include those involved in cell growth and development (LightGreen, LightYellow, LightCyan) and immune system processes (DarkRed and Brown); the modules most negatively associated with BW include those involved in lipid metabolic processes or mitochondrion function (Green, Blue, Cyan) and circadian rhythms (DarkTurquoise).

**Figure 3.**
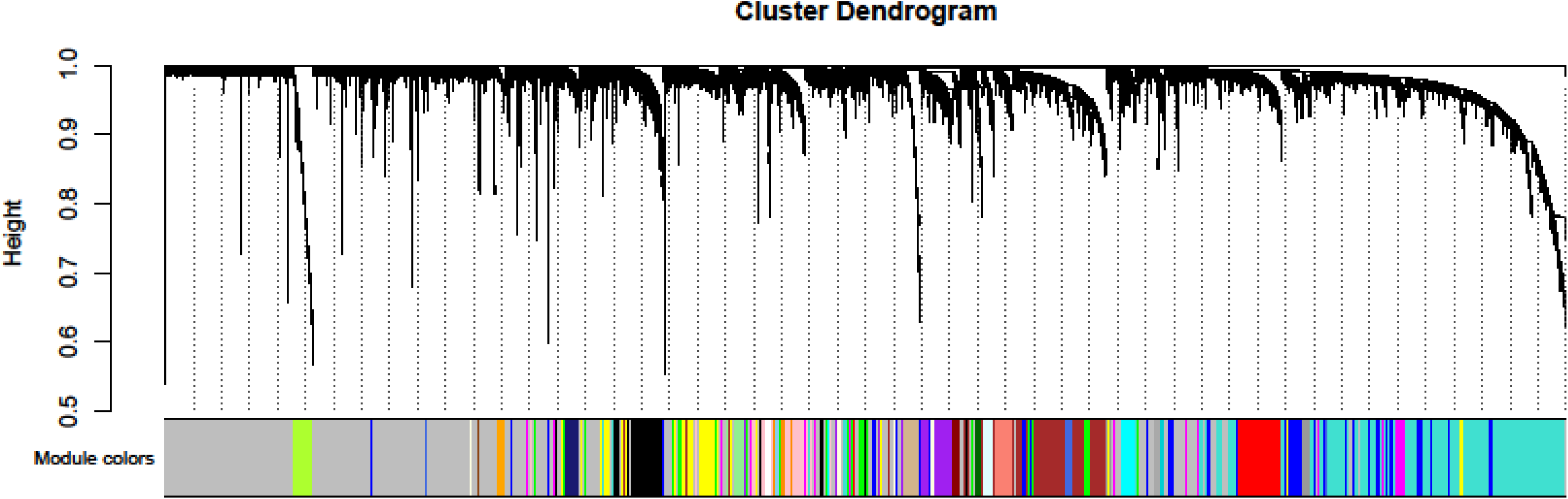
Cluster Dendogram for WGCNA analysis of HS rat adipose gene expression. Adipose gene expression was measured in 415 male HS rats. WGCNA analysis assigned 11,762 of 18,357 transcripts to 29 modules, ranging in size from 31 to 2,095 transcripts. Unassigned transcripts are shown in the ‘grey’ module.

**Figure 4.**
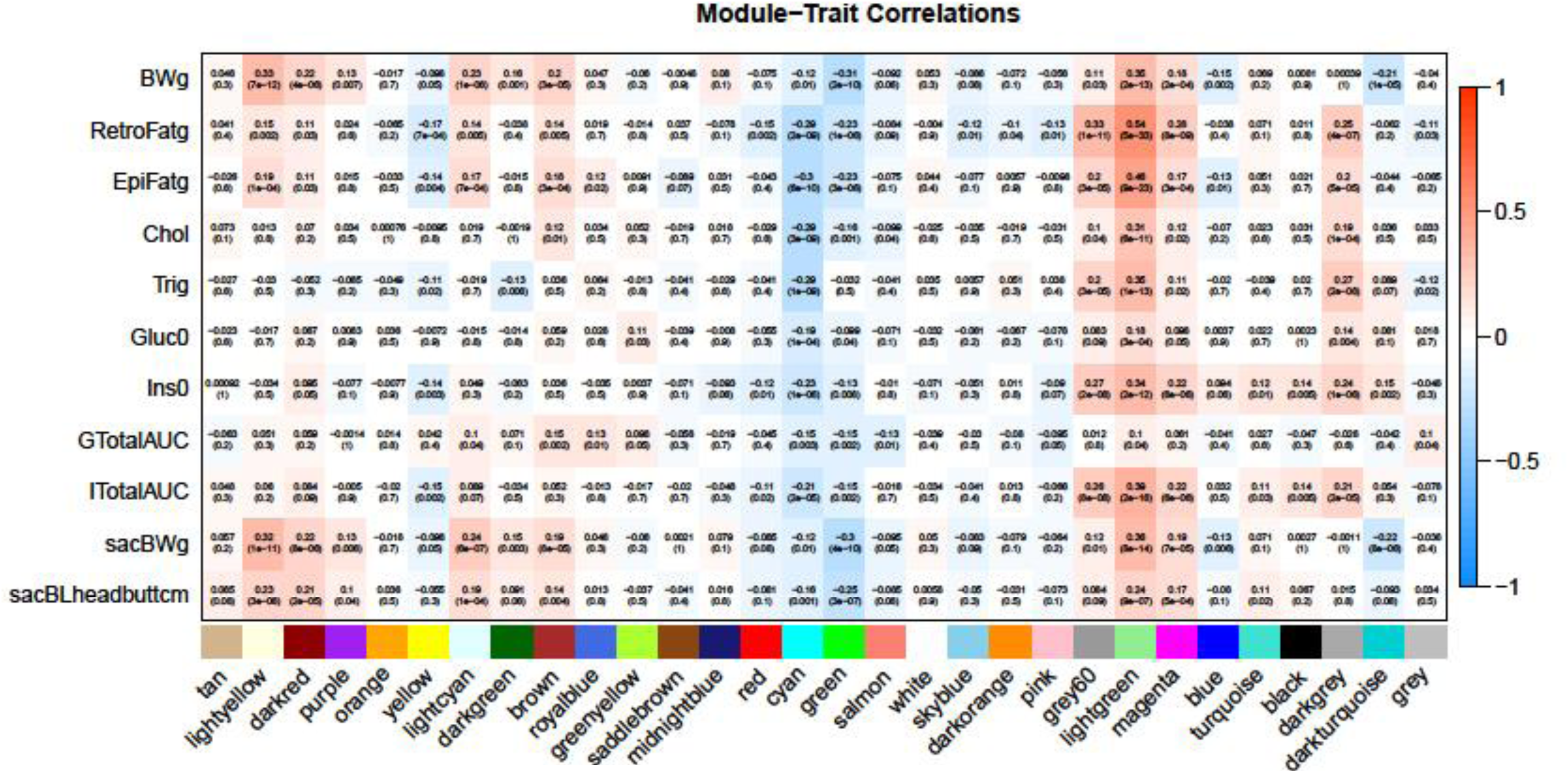
Correlations between WGCNA adipose tissue module eigengenes and metabolic phenotypes. Spearman correlations between measured metabolic phenotypes and module eigengenes from WGCNA analysis of adipose gene expression in 415 male HS rats. Eigengenes are the first principal component of gene expression for the genes included in each module. Several modules are significantly correlated with body weight (BWg) and other metabolic phenotypes. In particular, the ‘LightGreen’ and ‘LightYellow’ modules are most positively correlated with bodyweight, and the ‘Green’ module is most negatively correlated with bodyweight, while he ‘Cyan’ module is most negatively correlated with EpiFat and RetroFat. The strongest correlation of a module with any metabolic phenotype is between the ‘LightGreen’ module and RetroFatg. BWg – bodyweight (g); BLcm – body length (cm); EpiFatg – epididymal fat pad weight (g); RetroFatg – retroperitoneal fat pad weight (g); Ins0 – fasting insulin (ng/ml); InsAUC – insulin area under the curve after glucose challenge; Gluc0 – fasting glucose (mg/dL); GlucAUC – glucose area under the curve after glucose challenge; Chol – fasting total cholesterol (mg/dL); Trig – fasting triglycerides (mg/DL).

The LightGreen module was the most positively associated with BW (r = 0.35, p = 1.6x10-13) and was enriched for GO biological pathway (GO.BP) annotations including *positive regulation of pathway-restricted SMAD protein phosphorylation* (p=0.008; see **Table S7**), which is supported by canonical pathway enrichment analysis (**Fig. S4**). Hub genes in this module include *Adra2c*, *Myo1d*, and *Bace2* (highest intra-modular connectivity or IMConn, **Fig. 5** and **Table S3**), which are the most representative of the module (by correlation with the module eigengene, i.e. highest module membership). The LightGreen module has a strong positive correlation with several other metabolic traits including RetroFat, EpiFat, fasting cholesterol, triglycerides, insulin and insulin_AUC (**Fig. 4**). The LightYellow and LightCyan modules are also positively associated with BW (r = 0.33, p = 6.9 x 10-12 and r = 0.23, p = 1.5 x 10-06, respectively). These modules were enriched for GO annotations including *extracellular matrix organization* (p=.2 x 10-4 1; **Table S7**) and *cell cycle process (*p = 2.8 x 10-48). Together, these modules support a role of cell growth and development in adiposity.

**Figure 5.**
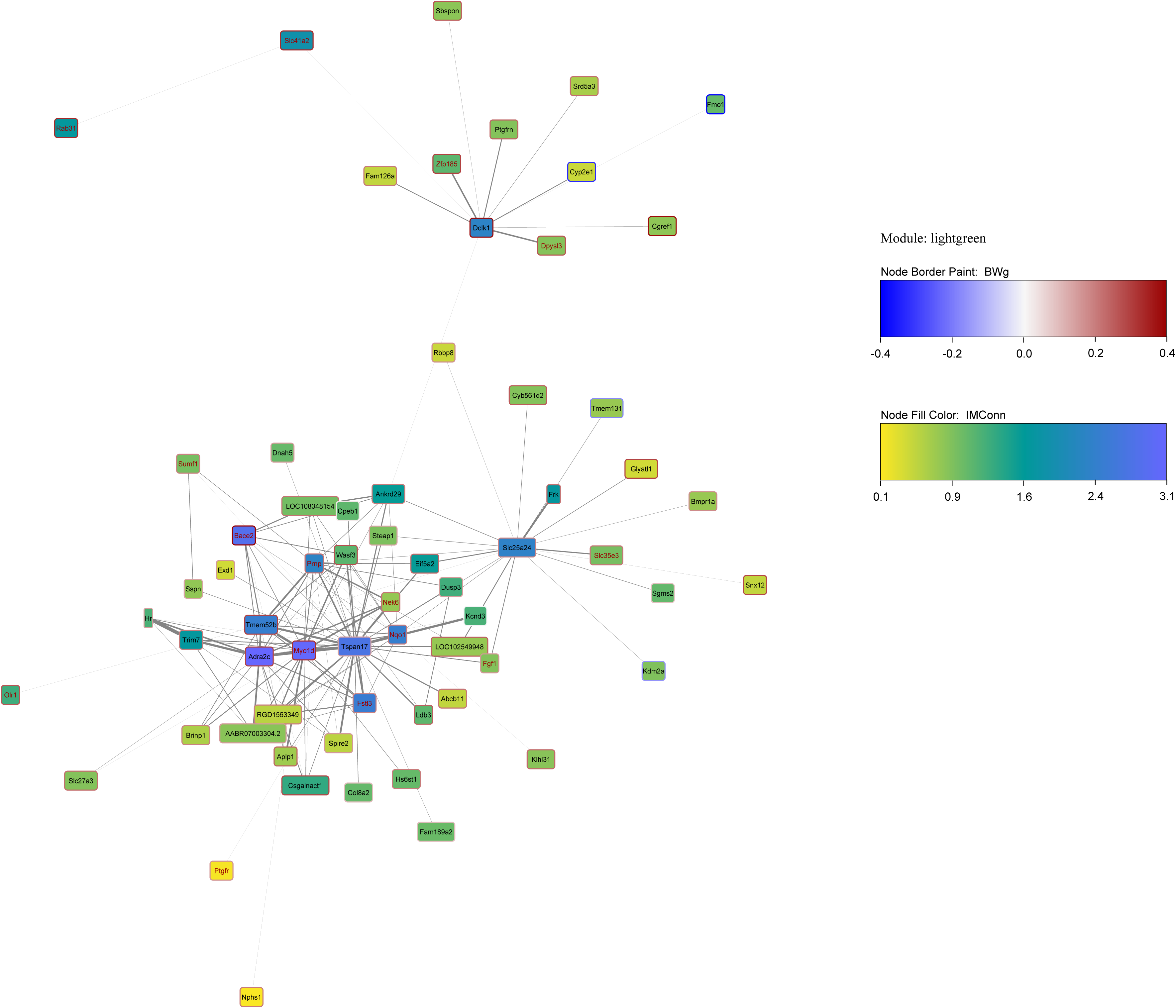
Network visualization of the LightGreen (SMAD Phosphorylation) module. Network visualization of the ‘LightGreen’ module from WGNCA analysis of adipose gene expression in HS rats. Edges are included if the adjacency between genes is greater than 0.03. Genes not connected to the main module networking have been removed. Edge width is scaled by adjaceny. The background of each node is colored by gene intramodular connectivity (IMConn). The border of each node is colored by the correlation between each gene and bodyweight (BWg). The label of each node is colored red if the gene is a consensus gene. Several genes are highly connected in the ‘LightGreen’ network, including *Adra2c, Bace2* and *Myo1d*, all of which are highly positively correlated with bodyweight, with *Bace2* and *Myo1d* being consensus genes.

The DarkRed and Brown modules are involved in immune system function and are strongly positively associated with BW (r= 0.22, p = 3.8 x 10-06 and r = 0.20, p= 3.2 x 1005, respectively). The DarkRed module is enriched for GO annotations including *complement activation*, and other immune-related terms, and the Brown module (a large network of 1138 genes) is enriched for GO annotations, including *immune system process* (p= 2.8 x 10-48), and many other immune-related terms (**Table S7**). KEGG pathway analysis for the DarkRed and Brown modules support a role in *complement and coagulation cascades* (p = 2.5 x 10-10), and *chemokine signaling pathway* (p = 1.7 x 10-13), respectively, as well as other immune-related terms (**Table S8**). Top hub genes for the DarkRed module include *Clec10a, Csf1r, C1qc, C1qa* and those for the Brown module include *Syk, Ptprc, Nckap1l, Arhgap30*, supporting a role of these modules in inflammatory and immune regulation mechanisms (**Table S3**).

Of those modules whose eigengenes most negatively correlated with BW, the Green module was enriched for *Cytokine Production* (p=4.7 x 10-05) using GO.BP (**Table 1** and **Table S7**), implicating immune system processes, which appears anti-intuitive. In contrast, analysis for KEGG pathways suggested enrichment for *Metabolic Processes* (p = 1.3 x 10-04). Upon closer investigation, 71 genes within this module have a strong positive correlation (r ≥ 0.2) with BW and were enriched for immune-inflammation related mechanisms (GO:BP 0002376∼immune system process, 26 genes, FDR-P= 3.03 x10^-5^), while 111 have a strong negative correlation (r ≤- 0.2) and were enriched for mitochondria (GO:CC 0005739∼mitochondrion, 21 genes, FDR-P=0.014), suggesting a role in inflammation mediated modulation of mitochondrial function. The Blue module (a large network of 1247 genes), is inversely correlated with BW (r= -0.15, p= 0.002), and is enriched for *mitochondrion organization* (p= 3.4 x 10-19) using GO.BP (**Table S7**) and *metabolic pathways* (p = 1.6 x 10-22) and *fatty acid metabolism* (2.9 x 10-08) using KEGG pathway analysis (**Table S8**).

Although only modestly associated with BW (r = -0.12, p = 0.014), the Cyan module is noteworthy for its strong negative correlation with several other traits including RetroFat (r = -0.29, p = 2.4 x 10-09), EpiFat (r = -0.30, p = 6.3 x 10-10), fasting insulin (r = -0.23, p = 1.3 x 10-6) and fasting cholesterol (r = -0.29, p = 3.2 x 10-09) and triglycerides (r = -0.29, p = 1.2 x 10-09) (**Table 1** and **Fig. 4**). The Cyan module is enriched for GO annotations such as *regulation of neurotransmitter levels* (p = 0.004) and *lipid metabolic process* (p = 0.008; see **Table S7**), with *thyroid hormone signaling pathway* being significant in KEGG pathway analysis (p = 1.7 x 10-04; **Table S8**). Hub genes in the Cyan module include *Psat1* and *Acvr1c* (**Fig. 6**). Together, these modules implicate an inverse relationship between lipid metabolic processes and mitochondrion function with adiposity.

**Figure 6.**
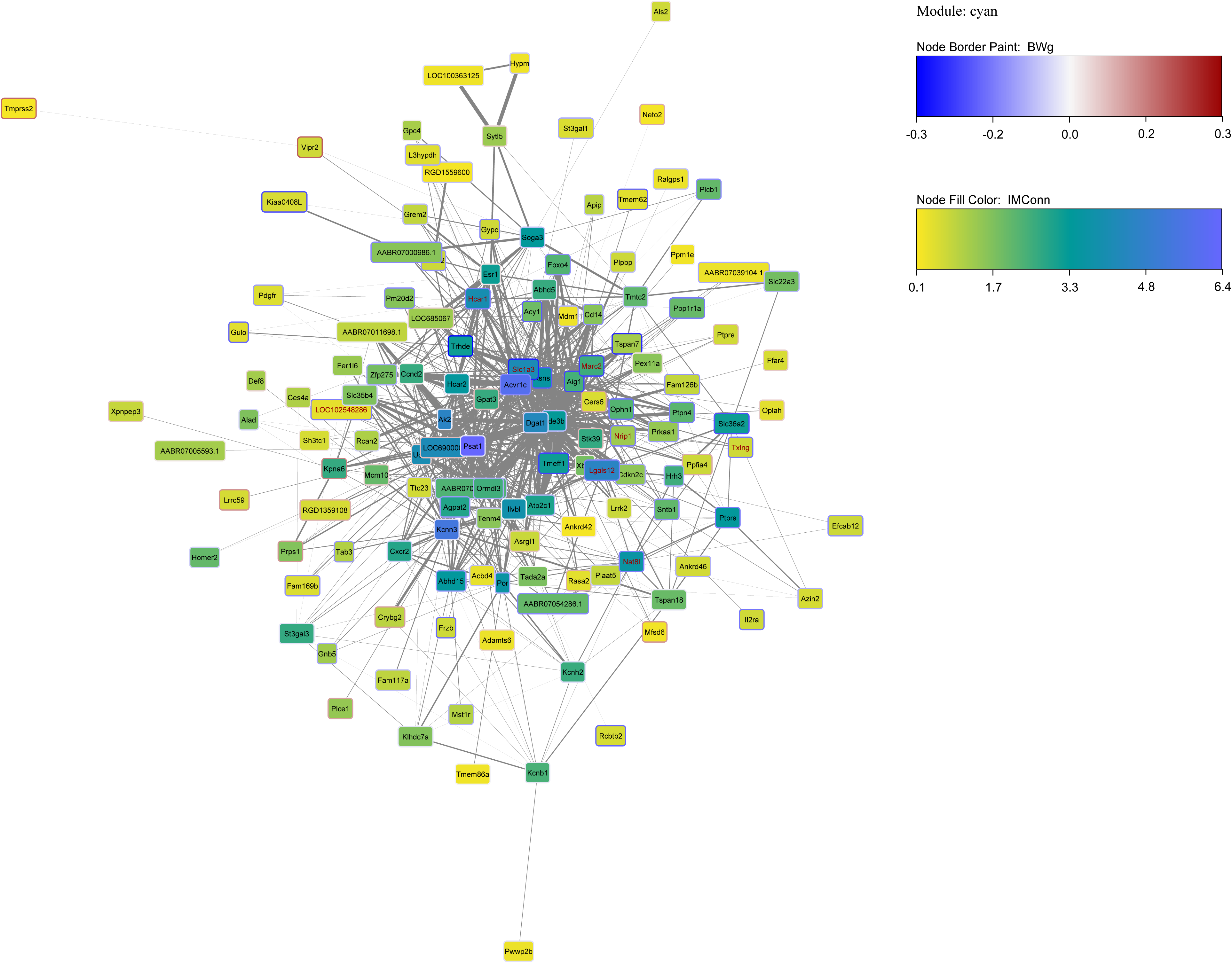
Network visualization of the Cyan (Regulation of neurotransmitter levels and lipid metabolic process) module. Network visualization of the ‘Cyan’ module from WGNCA analysis of adipose gene expression in HS rats. Edges are included if the adjacency between genes is greater than 0.03. Genes not connected to the main module networking have been removed. Edge width is scaled by adjaceny. The background of each node is colored by gene intramodular connectivity (IMConn). The border of each node is colored by the correlation between each gene and bodyweight (BWg). The label of each node is colored red if the gene is a consensus gene. Several genes are highly connected in the ‘Cyan’ network, including *Acvr1c, Slc1a3*, *Lgals12,* and *Hcar1,* all of which are negatively correlated with bodyweight and with the latter three being consensus genes and *Hcar1* showing evidence of mediation.

The DarkTurquoise module is also highly negatively correlated with BW and is enriched for *Circadian Rhythm* (p=0.007) which is supported by KEGG pathway analysis (p=4.4 x 10-04) with hub genes including *Prex2* and *Gpd1l* (**Tables S3, S7, S8**).

### BW associated transcripts in rat adipose often associate with BMI in human adipose

We identified a sub-set of “consensus” genes whose expression levels were independently and significantly associated with BW in HS rat adipose and BMI in human adipose (FDR = 1%) with the same direction of effect in both AAGMEx (662/4669 BMI associated genes) and METSIM (896/6058 BMI associated genes) cohorts. There is substantial overlap between the three cohorts as shown in **Table S9**. In total, 554 consensus genes were correlated in all three data-sets with the same direction of effect -- an overlap between studies that is significantly higher than expected by chance (chi-squared p < 0.00001). Ranking genes by their Fisher combined p-value across all three datasets, the most significantly associated genes were *Spx, Hadh, Slc27a2, Cmtm3, Ccn5, Htra1, Slc4a4, Letmd1, Echdc3, Crls1* (**Table S10**).

These 554 consensus genes are enriched in 8 of the 29 rat adipose co-expression modules identified by WGCNA (**Table 1**), comprising 341 of the 554 (61.5%) consensus genes. These 8 modules include those whose module eigengene was highly correlated with BW in the rat as described above and in **Table 1**.

### Consensus genes may have a causal genetic effect on rat BW via expression levels

Of the 554 consensus genes, 548 had SNPs within 1 Mb of the gene in the rat, 149 of which also had a cis-eQTL (**Table S11**). We applied mediation analysis to those 149 genes and identified 19 whose adipose expression may mediate a genetic effect on BW in rat (**Table 2**). Of these, 14 show evidence for complete mediation (ie, the effect of the cis-eQTL’s top SNP on BW is entirely mediated through expression of the gene) and 5 have evidence for partial mediation. The gene *St14*, a known cancer gene (36), has the most significant association (p=0.002) between BW and its lead cis-eQTL SNP among all candidate mediators and is a complete mediator of this association. The gene *Htra1*, a negative regulator of adipogenesis (37), was among the most significant consensus genes by Fisher combined p-value (across all three datasets), and its lead cis-eQTL SNP is nominally associated (p<0.05) with BW. Two mediators act as hub genes: *Hcar1* in the Cyan module and *Ms4a6a* in the Green module. Three of the mediators play a role in inflammation (*Rgs1, Vcam1, St3gal5*), and several genes play a role in adipogenesis (*Fzd7, Tmem120b, Atl2, Htra1, St3gal5*). Other mediators are involved in lipolysis (*Hcar1*), fatty acid oxidation (*Mlycd*), cell-cell interactions (*Cpxm2*), oxidative stress (*Cisd2*) or weight maintenance (*Pros1*). Seven genes have no previously known role in obesity (*St14, Ms4a6a, Ostc, Ung, Tspan33, Tigd2, Mocos;* **Table 2**). Using PhenomeXcan, we found that six of the 19 genes (*Fzd7, Hcar1, Ms4a6a, Htra1, Tigd2, Cpxm2*) exhibit potential causal relationships with adiposity related traits in human, with three genes (*Cisd2, Ms4a6a, St3gal5*) showing associations with monocyte count or percentage, with the same direction of effect as seen in the rat (**Table 2;** details are in **Table S12**).

**Table 2.** Genes identified through mediation analysis that may play a causal role in obesity

| Gene Name | Gene Description [RGD Acc#] | SNP basepair | Model 1: (y ~ x') | Model 2: (m ~ x') | Model 3: (y ~ m' + x) | Model 4: (y ~ m + x') | Support (PMID) | Entrezgene ID | Module/ Hub (y/n) | Association in Phenome Xcan |
| --- | --- | --- | --- | --- | --- | --- | --- | --- | --- | --- |
| *St14* | suppression of tumorigenicity 14 [Acc:69288] | chr8<br>32379011 | 0.002045 | 3.00E-13 | 5.98E-05 | 0.156456 | none | 114093 | Brown/No | No |
| *Fzd7* | frizzled class receptor 7 [Acc:2321905] | chr9<br>67275996 | 0.003163 | 2.06E-07 | 0.014260 | 0.027514 | 25871514<br>23861788<br>21354690 | 100360552 | Green/No | Yes |
| *Hcar1* | hydroxycarboxylic acid receptor 1 [Acc:1586364] | chr12<br>38233140 | 0.003572 | 2.44E-05 | 0.020101 | 0.017515 | 33284607<br>31999787<br>22842580<br>20374963<br>19633298 | 689936 | Cyan/Yes | Yes |
| *Cisd2* | CDGSH iron sulfur domain 2 [Acc:1566242] | chr2<br>240335511 | 0.004145 | 5.69E-08 | 0.000962 | 0.056948 | 29556009<br>25448035<br>33610659 | 295457 | Light Green/No | Yes |
| *Ms4a6a* | membrane spanning 4-domains A6A [Acc:1305800] | chr1<br>227335280 | 0.009635 | 7.03E-07 | 0.001267 | 0.079024 | none | 361735 | Green/Yes | Yes |
| *Ostc* | oligosaccharyltransferase complex non-catalytic subunit [Acc:1560708] | chr2<br>235860726 | 0.015812 | 1.24E-06 | 3.47E-05 | 0.000744 | none | 362040 | Turquoise/No | No |
| *Rgs1* | regulator of G-protein signaling 1 [Acc:3561] | chr13<br>61138665 | 0.016762 | 3.16E-11 | 0.000989 | 0.219277 | 25744306<br>27561966 | 54289 | Brown/No | No |
| *Ung* | uracil-DNA glycosylase [Acc:1307200] | chr12<br>48223578 | 0.017222 | 3.04E-06 | 0.007930 | 0.091762 | none | 304577 | Light Yellow/No | No |
| *Tmem120b* | transmembrane protein 120B [Acc:1584031] | chr12<br>39084074 | 0.017389 | 1.64E-06 | 0.018515 | 0.076418 | 26024229 | 690137 | Brown/No | No |
| *Atl2* | atlastin GTPase 2 [Acc:1305125] | chr6<br>2465360 | 0.022600 | 1.41E-05 | 1.08E-07 | 0.251464 | 32349335 | 298757 | Blue/No | No |
| *Tspan33* | tetraspanin 33<br>[Acc:1560915] | chr4<br>57238350 | 0.037508 | 3.08E-12 | 0.003185 | 0.327096 | none | 500065 | none | No |
| *Htra1* | HtrA serine<br>peptidase 1<br>[Acc:69235] | chr1<br>201245066 | 0.038328 | 5.86E-16 | 1.78E-05 | 0.819508 | 26864869<br>31517621 | 65164 | Turquoise<br>/No | Yes |
| *Vcam1* | vascular cell<br>adhesion<br>molecule 1<br>[Acc:3952] | chr2<br>219251193 | 0.040169 | 2.69E-09 | 5.11E-08 | 0.697793 | 253 pubmed<br>refs | 25361 | Green/No | No |
| *Mlycd* | malonyl-CoA<br>decarboxylase<br>[Acc:620234] | chr19<br>53006509 | 0.041178 | 0.000129 | 0.000585 | 0.170482 | 25016691<br>30481042 | 85239 | Blue/No | No |
| *Tigd2* | tigger<br>transposable<br>element derived<br>2 [Acc:1559612] | chr4<br>90047090 | 0.047026 | 7.04E-15 | 0.009026 | 0.379197 | none | 100912924 | none | Yes |
| *Mocos* | molybdenum<br>cofactor<br>sulfurase<br>[Acc:1308496] | chr18<br>15927958 | 0.047739 | 9.96E-15 | 1.41E-05 | 0.000372 | none | 361300 | Dark<br>Turquoise<br>/No | No |
| *Cpxm2* | carboxypeptidase<br>X (M14 family),<br>member 2<br>[Acc:1310955] | chr1<br>204245737 | 0.049090 | 2.78E-37 | 7.60E-08 | 0.112586 | 24685769 | 293566 | none | Yes |
| *Pros1* | protein S<br>[Acc:620971] | chr7<br>1206510 | 0.049665 | 2.25E-10 | 0.001241 | 0.003590 | 21957032 | 81750 | Turquoise<br>/No | No |
| *St3gal5* | ST3 beta-<br>galactoside<br>alpha-2,3-sialyltransferase<br>5 [Acc:620875] | chr4<br>99943651 | 0.049891 | 1.38E-10 | 0.001547 | 0.373870 | 26499445<br>34125497 | 83505 | none | Yes |

Genes highlighted in gray under model 4 show evidence for partial mediation. All others show evidence for complete mediation.

## DISCUSSION

We found functional concordance between BW associated adipose tissue transcripts in the HS rat and their human BMI-associated orthologues. This work replicates previous findings in human (38) and mouse (20), and establishes the utility of the HS rat, an outbred model previously used for the genetic mapping of complex traits (8-11, 34, 39), as a model to understand the systems genetics of human obesity. Specifically, we observed many more orthologous genes with the same pattern of correlation with rat BW or human BMI than would be expected by chance, confirming that the systems genetics of these traits are at least partially evolutionarily conserved between humans and rodents despite 90- to 100 million years of evolutionary divergence. The human/rat concordant genes cluster into rat gene networks involved in cell growth and development and immune system function (positive associations) and lipid metabolic processes and circadian rhythms (negative associations). Although network analysis alone cannot determine whether modules are causal or reactive to the obesity state, by integrating genotype data, we identified 19 genes whose expression levels are genetically regulated and which appear to mediate the genetic effects on BW, suggesting a causal role for these genes. Although several of these genes have previously been implicated in obesity, many are novel, opening up new targets to understand the mechanistic underpinnings of obesity.

Several BW-associated co-expression modules are involved in immune system function (Brown, DarkRed, Green). This is not surprising as obesity is known to activate the immune system (40, 41) and previous work has shown similar findings in both human (38) and mouse adipose tissue (20). The current work specifically suggests involvement of genes involved in Fc-gamma receptor-mediated phagocytosis, the chemokine signaling pathway and complement activation. For example, the Brown module was enriched for the chemokine signaling pathway and cytokine-cytokine receptor interaction. Among the top hub genes in this module, Rho GTPase Activating Protein 30 (*Arhgap30)* is a hub gene in human adipose tissue co-expression modules associated with triglyceride level (42), as well as BMI and insulin resistance (21). The DarkRed module was enriched for complement and coagulation cascades and top hub genes in this module include *complement C1q C chain and A chain* (*C1qc* and *C1qa*), as well as *C-type lectin domain containing 10A* (*Clec10a*) and *Colony stimulating factor 1 receptor* (*Csf1r*). *Clec10a* has previously been identified as playing a causal role in insulin resistance in African Americans via adipose tissue expression levels (21), while blockade of *Csf1r* depletes macrophages and prevents fat storage and adipocyte hypertrophy in high-fat diet-fed and hyperphagic mice (43). The immune modules were also strongly enriched for consensus genes, implying conservation of obesity-associated immune-inflammation pathways between rats and humans.

Other BW-associated modules that are enriched for consensus genes are involved in cellular growth and development including the LightGreen (*regulation of pathway-restricted SMAD protein phosphorylation*) and LightYellow (*extracellular matrix (ECM) organization*) modules. Phosphorylation of SMAD proteins leads to cell cycle inhibition (44). In addition to their roles in cell proliferation and differentiation, SMAD proteins are involved in extracellular matrix (ECM) remodeling and immune function (45). Excessive ECM protein deposition, followed by fibrosis in adipose tissue, is considered a pathological consequence of long-term obesity and has been observed in rodents and humans (40). Consistent with our findings, adipose tissue TGFbeta signaling is upregulated in obesity in mice (46) and humans (47) and these changes are associated with decreased adipogenesis (46) as well as collagen expression and fibrosis (47). In addition, SMAD3 knock-out mice exhibit decreased white adipose tissue mass with improved glucose tolerance and insulin sensitivity (48, 49). Thus, increased TGFbeta signaling and SMAD phosphorylation in obesity likely decreases cell differentiation and increases fibrosis.

Consensus enriched modules that were most negatively associated with BW are involved in lipid metabolic processes (Cyan), mitochondria organization (Green, Blue) and circadian rhythms (DarkTurquoise). Each of these processes are often disrupted in obesity and play an important role in disease process (50, 51). Specifically, the Cyan module is enriched for GO terms including regulation of neurotransmitters and lipid catabolic processes. Catecholamines are activated during fasting or stress and lead to beta-adrenergic signaling in adipose tissue to mobilize stored energy through lipolysis and/or thermogenesis. Although basal lipolysis has been shown to be upregulated in obesity (52), catecholamine stimulated lipolysis is decreased, and may play a role in further exacerbating the obese condition (53). Our work shows a negative relationship between obesity and catecholamine signaling and/or lipolysis, further supporting this finding. *Hydroxycarboxylic Acid Receptor 1 (Hcar1)*, a hub gene in this module and one of the genes identified through mediation analysis, is regulated by PPARgamma (54) to mediate anti-lypolitic events (55). Our data indicates that increased expression of *Hcar1* drives up lipolysis resulting in decreased fat pad size.

In addition to identifying functionally relevant pathways for obesity between the HS rat and humans, we identified consensus genes that may play a causal role in obesity via genetic variants that drive transcript expression. These include genes involved in adipogenesis (*Fzd7, Tmem120b, Atl2, Htra1, St3gal5*) and inflammation (*Rgs1, Vcam1, St3gal5*), among others (see **Table 2**). Although inflammation is generally thought to be responsive to obesity and over-nutrition, this work, along with findings from human GWAS (15), indicates that inflammation may also play a causal role. We also identified several genes with no previously known connections to obesity (*St14, Ms4a6a, Ostc, Ung, Tspan33, Tigd2, Mocos*), opening up new avenues of research to explore novel mechanistic underpinnings of obesity. For example, *St14* is a serine protease involved in cancer metastasis (36) that has not previously been linked to obesity. Using PhenomeXcan (35), we found that eight of the 19 genes exhibit a potential causal gene-trait relationship with adiposity or immune phenotypes in human, including *Ms4a6a* and *Tigd2,* two genes with no known previous role in adiposity. *Ms4a6a* is also a hub gene in the Green module, is expressed in the hematopoietic system and has previously been associated with Alzheimer’s disease (56), making it a particularly attractive candidate for obesity. Importantly, because environmental conditions likely differ significantly between rat (chow, non-obesogenic diet) and human (uncontrolled diet, likely Western, obesogenic), we do not believe that these findings rule out the possibility that these other genes also play a role in human adiposity. A potential role of the 19 mediator genes in human obesity is supported by the fact that each of the mediators have an orthologous cis-eQTL in the METSIM human cohort.

In summary, we have shown that many BW associated genes, as well as their associated networks and pathways, in the HS rat are conserved in humans, indicating similar pathways regulate obesity in both species. We identified a sub-set of genes that may play a causal role in obesity, and these genes encompass mechanisms involved in adipocyte differentiation and inflammation, with several genes being novel. These findings support the HS rat as a model to study the systems genetics of obesity and identifies novel biological targets for future functional testing.

## ACKNOWLEDGMENTS

Funding: R01 DK 106386 (LSW), R01 DK120667 (LSW), R01 DK090111 (SKD), R01 DK118243 (SKD) R35 GM127000 (WV) The authors thank the METSIM study investigators for publicly sharing their data and summary statistics.

Authors declare no conflicts of interest for this work.

